# Choline Transporter in α/β core neurons of *Drosophila* mushroom body non-canonically regulates pupal eclosion and maintains neuromuscular junction integrity

**DOI:** 10.1101/380519

**Authors:** Runa Hamid, Nikhil Hajirnis, Shikha Kushwaha, Sadaf Saleem, Vimlesh Kumar, Rakesh K Mishra

## Abstract

Insect mushroom bodies (MB) have an ensemble of synaptic connections well-studied for their role in experience-dependent learning and several higher cognitive functions. MB requires neurotransmission for an efficient flow of information across synapses with the different flexibility to meet the demand of the dynamically changing environment of an insect. Neurotransmitter transporters coordinate appropriate changes for an efficient neurotransmission at the synapse. Till date, there is no transporter reported for any of the previously known neurotransmitters in the intrinsic neurons of MB. In this study, we report a highly enriched expression of Choline Transporter (ChT) in *Drosophila* MB. We demonstrate that knockdown of ChT in a sub-type of MB neurons called α/β core (α/βc) neurons leads to eclosion failure, peristaltic defect in larvae, and altered NMJ phenotype. These defects were neither observed on knockdown of proteins of the cholinergic locus in α/βc neurons nor by knockdown of ChT in cholinergic neurons. Thus, our study provides insights into non-canonical roles of ChT in MB.

## Introduction

Acetylcholine (ACh) is essential for higher cognitive functions and in many of the developmental events of CNS. The components of ACh metabolic cycle namely, Choline acetyltransferase (ChAT), vesicular acetylcholine transporter (VAChT), Acetylcholine esterase (AChE) and Choline transporter (ChT) work in synchronization with each other to bring efficient neurotransmission at cholinergic synapses. Timely removal of ACh from synaptic cleft is a key step of synaptic transmission mediated by ChT. For resynthesis of ACh, ChT transports choline into the presynaptic terminal which is produced by the enzymatic action of Acetylcholine esterase (AChE) at the synapse. There are few, but intriguing evidence, that associate ChT with neuromuscular dysfunction, congenital myasthenic syndrome, severe neurodevelopmental delay and brain atrophy (Barwick et al., 2012; Wang et al., 2017). ChT mutants demonstrate motor activity defects in C. *elegans* (Matthies et al., 2006). Alzheimer’s disease has also been associated with altered levels of ChT in cholinergic neurons (Bissette et al., 1996; Pascual et al., 1991). The cognitive significance of ChT has also been indicated in rodents performing various tasks (Durkin, 1994; Toumane et al., 1989; Wenk et al., 1984). Although sparse, all the previous studies bring into focus a highly important role of ChT in cholinergic nerve terminals.

A recent study demonstrates that intrinsic neurons of *Drosophila* mushroom bodies (MB) are principally cholinergic and express ChAT and VAChT. (Barnstedt et al., 2016). This contrasts with the previous findings that show components of the cholinergic cycle are absent from MB (Gorczyca and Hall, 1987; Yasuyama et al., 1995). For an efficient cholinergic neurotransmission in these structures, all the components of the cholinergic cycle should be present. This knowledge is still obscure. It remains unclear whether: (a) all the intrinsic cells of MB are cholinergic in nature or (b) different cells represent a different type of neurochemical. A specific subset of MB intrinsic neurons called α/β core (α/βc) neurons transiently uses Glutamate as a neurotransmitter at the time of eclosion (Sinakevitch et al., 2010). Studies also show the presence of aspartate and taurine in a limited number of intrinsic neurons of MB (Sinakevitch et al., 2001). However, Glutamate transporter, vGLUT, is absent from core neurons or other intrinsic neurons of MB (Daniels et al., 2008). Transporters for GABA (DvGAT) and monoamines (DvMAT) are also absent in intrinsic neurons of MB (Chang et al., 2006; Fei et al., 2010). Taken together, different studies describe different neurotransmitter in intrinsic neurons of MB but the transporters for the known neurotransmitters are absent from these cells. *Drosophila* portabella gene has been previously reported as a putative transporter in MB, but no definitive neurotransmitter has been defined for it (Brooks et al., 2011). In the absence of any transporter, it is not clear how neurotransmitter release and consequently different synaptic strengthening is achieved which might underlie cognitive and developmental events in this structure.

Here, we report for the first time, the presence of ChT in MB and demonstrate a distinct localization of endogenous ChT in all the major lobes of MB. Knockdown of ChT specifically in α/β core (α/βc) neurons leads to severe eclosion failure without affecting any larval or pupal development. We also report peristalsis defect and altered NMJ phenotype in these animals. All the three phenotypes: eclosion failure, peristaltic defect, and altered NMJ phenotype are rescued to a significant amount by transgenic over-expression of ChT in α/βc neurons. Furthermore, we demonstrate that the function of ChT in α/βc neurons is independent of the cholinergic pathway. Our results suggest that the role of ChT to transport choline for ACh synthesis might not be exclusive, at least in α/βc neurons. Together, our study reveals a new marker for the *Drosophila* MB and suggests its specific role in eclosion and maintenance of NMJ integrity. Defective NMJ due to knockdown of ChT might be the underlying cause of eclosion failure. In view of any transporter being absent in MB, our findings have broad implications in understanding the functioning of the neural circuits in MB – a region that controls animal behavior and higher cognitive functions.

## Results

### Choline transporter is enriched in *Drosophila* mushroom body

ChT is a phylogenetically conserved protein. It was first identified in *C.elegans* followed by rat, mouse, humans and other species (Apparsundaram et al., 2001; Apparsundaram et al., 2000; O’Regan et al., 2000; Okuda et al., 2000; Wang et al., 2001). It has a high binding affinity for choline (Km ~ 1-5 μM) in the nervous system (Kuhar and Murrin, 1978; Lockman and Allen, 2002). *Drosophila* genome also has an annotated ChT homolog, *CG7708.* To study the function of ChT in CNS, we generated a polyclonal antibody against the 125 amino acid long C-terminal of ChT protein. Immunostaining of ventral ganglia with pre-immune sera did not show any immunoreactivity whereas affinity purified anti-ChT serum showed predominant immunoreactivity in the neuropil of ventral nerve cord (VNC) of the third instar larval brain (Fig.S1 A-F). This suggests an enrichment of the endogenous ChT protein at the neuropilar synapses. We assessed the specificity of this antibody for the endogenous ChT protein. Immunostaining of larval ventral ganglion of *Elav^Cl55^GAL4* driven *ChT* RNAi (ChT^RNAi^) showed a significant reduction of ChT immunoreactivity at the central synapses of VNC as compared to control while costaining with anti-ChAT showed no reduction in immunoreactivity of ChAT protein (Fig. S1 G-L). In addition, to determine if the ChT is localized at cholinergic synapses, we assessed colocalization of ChT protein with canonical proteins of the cholinergic cycle, ChAT, and VAChT of the cholinergic locus. By immunostaining of 3^rd^ instar larval VNC, we observed an extensive colocalization of ChT with both ChAT and VAChT in the neuropilar areas (Fig.S2 A-F). ChT also colocalizes with ChAT in other cholinergic synaptic rich regions of the brain like Bolwig nerve (Fig.S2 G-I) and antennal lobes (AL) (Fig S2. J-L). Together, these data show that ChT antibody specifically recognizes endogenous ChT protein at cholinergic synapses of VNC.

In *Drosophila* central brain, we observed that ChT staining was pronounced at the neuropil of the larval central brain and sub-oesophageal ganglia (SOG). It colocalizes extensively with ChAT (Fig. 1A). Strikingly, we observed expression of ChT but not ChAT in the MB of the larval brain (Fig. 1A, A’). This pattern of expression was maintained post-metamorphosis in adult brain as well (Fig. 1B, B’). During development of the adult brain, the first lobe of MB formed is the ϒ lobe which grows until mid-third instar larval stage. The next is, α’/β’ which continues to form till puparium formation. Lastly, α/β lobes are formed from the puparium stage until adult eclosion (Lee et al., 1999). Immunostaining with anti-Disc large (anti-Dlg) and costaining with anti-ChT showed an immense localization of ChT in all the three lobes as well as in spur region of MB (Fig. 1C, D, E, E’). Altogether, these data suggest that ChT is explicitly expressed in all the lobes of *Drosophila* MB. Given the importance of MB in insect behavior, the ubiquitous expression of ChT in MB is intriguing, indicating ChT to be a critical protein for MB physiology and functioning.

**Figure 1:**
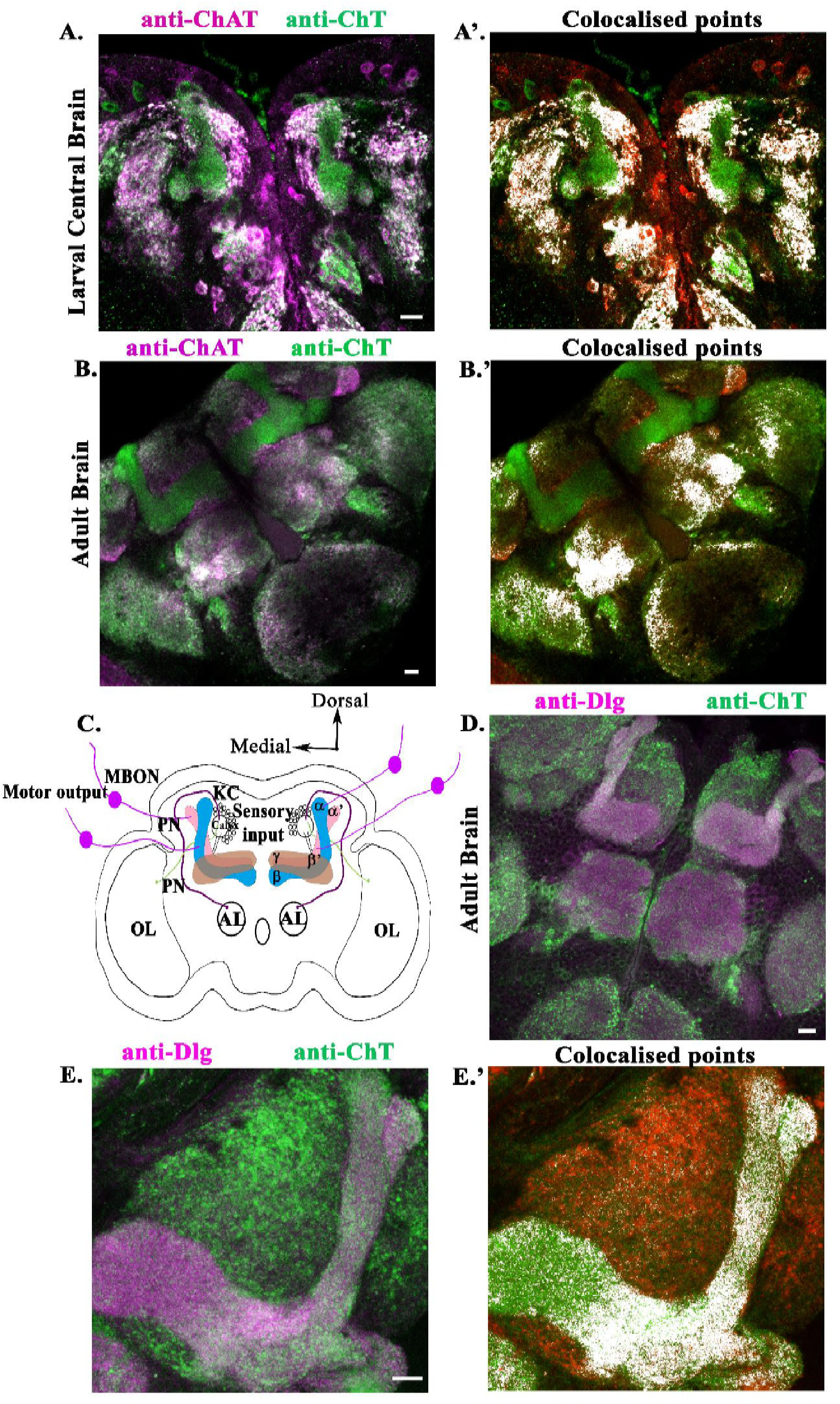
ChΓ is expressed in MB of larval as well as adult Drosophila brain. *(A)* left panel, shows co-immunostaining of ChT and ChAT in the central brain of 3^rd^ instar larva, colocalized regions are shown as white, (A’) right panel represents the same image processed by image J showing the colocalized pixels as white. *(B)* shows co-immunostaining of ChT and ChAT in the dissected adult fly brain (colocalized regions, white). The L-shaped structures are MB. (B’) shows colocalized pixels as white. *(C),* schematics of the fly adult brain showing MB and different neural processes in and out of it. MB circuitry is formed by a posterior cluster of about 2200-2500 Kenyon cells (KC) shown here as empty small circles. KC extend their dendritic processes to form calyx. Projection neurons (PN) from different sensory glomeruli project their axons into the MB calyx. Shown here are antennal lobe (AL) and optic lobe (OL) and representative PN coming out from them as magenta and green respectively. Axons of KCs form fasciculated axonal tract called peduncle which branches into lobes, bifurcating dorsally to form α/α’ and medially β/β’ (shown as pink and blue lobes). A single ϒ lobe which is continuous with heel wraps β’ lobe (shown as brown lobe). The MB lobes synapse with dendrites of Mushroom body output neurons (MBON) which provide the motor input and is the only output of MB (purple solid circle). *(D)* shows the co-immunostaining of Dlg and ChT protein in the adult *Drosophila* brain. (E) shows digitally zoomed image of Dlg and ChT co-immunostained single MB of the adult brain (left panel) and processed colocalized image (E’). All immunostained images are pseudocolored z-stack confocal images which are representative of 5-7 brains. Scale bar, 50 μm.

### ChT function in α/β core neurons of the mushroom body is required for eclosion

To investigate the functional relevance of ChT in MB, we used RNAi transgene of *ChT* (ChT^RNAi^) to cause a reduction in the expression of *ChT* mRNA levels. For this, we used GAL4 drivers specific for expression in different lobes of MB (Aso et al., 2009). The expression pattern of the driver lines was checked by driving expression of *mCD8GFP* with GAL4 and immunostaining of adult brain with anti-ChT. *201Y*GAL4 showed specific expression in densely packed fibers of α/β core (α/βc) in the center and ϒ lobe (Fig. 2A). Knockdown of ChT in these neurons showed eclosion failure in more than 90% of progeny as compared to the control (Fig.2C and Table 1). The co-staining with anti-ChT and anti-Dlg showed that the anatomy of MB is intact and there is no apparent developmental defect in the overall morphology of the brain due to knock down of ChT (Fig. 2B). We found that *201Y*GAL4>ChT^RNAi^ flies develop normally throughout the larval and pupal stages (Fig. 2D). However, from these observations, we cannot rule out the possibility of any developmental defect in the establishment of neural circuits, caused due to the reduction of ChT. Indeed, the flies are alive till more than P14 stage (Video1). Some of the flies were also unable to push themselves out of the pupal case consequently leading to the death of the flies (Video 2). 10% flies escape, display severe flight and motor defects and die within a day or two (Fig. 2C and Table 1). The escapies also show abnormal abdominal melanization and wing expansion failure (data not shown). To confirm if the eclosion failure is due to knock down of ChT in α/βc neurons, the fly was removed from the pupal case and brain was dissected out. Immunostaining of dissected adult brain of *201Y*GAL4>ChT^RNAi^, with anti-ChT, showed significant reduction of ChT intensity normalized to the ChT intensity in the neuropilar areas outside MB (Control 1.02±0.09, N=12; *201Y*GAL4>ChT^RNAi^ 0.48±0.05, N=12; Fig. 2E and 2G). Transgenic overexpression of UAS-ChT in MB by *201Y*GAL4 significantly restored ChT levels in MB (*201Y*GAL4/ChT^RNAi^;UAS-ChT/UAS-ChT, 0.88±0.05, N=18, Fig. 2F and G). We also observed rescue of eclosion failure to a significant level by expression of a single copy of UAS-*ChT* transgene and a double copy of UAS-ChT transgene (Fig. 2H and Table 1). Taken together these data suggest a specific role of ChT in α/βc neurons to regulate eclosion of adult *Drosophila* flies from the pupal case.

**Figure 2:**
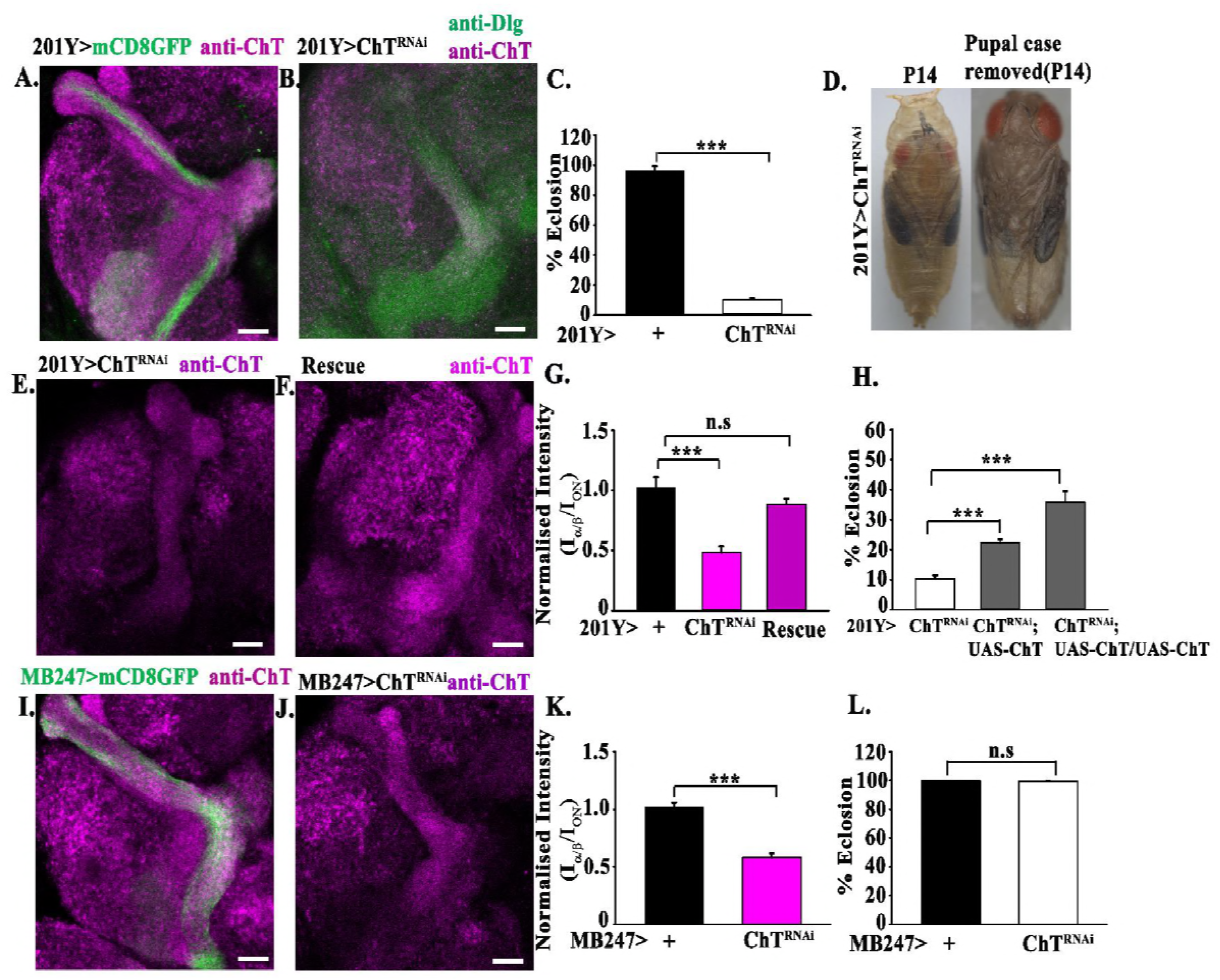
Downregulation of ChT in α/βc neurons lead to eclosion failure which is rescued by transgenic overexpression of ChT. (A) Shows mCD8GFP driven by 201YGAL4, marks α/βc neurons and ϒ lobe (green) and co-stained by anti-ChT (magenta) in MB lobe. (B) knockdown of ChT by ChT^RNAi^ is driven by 201YGAL4 and coimmunostained with anti-ChT (purple) and anti-Dlg (green). (C) shows percent eclosion failure by knockdown of ChT in 201Y>ChT^RNAi^ as compared to 201Y>+. (D) shows development in P14 staged undissected pupa (left) and fly dissected out from the pupal case (right) of 201Y>ChT^RNAi^ genotype. (E) shows anti-ChT immunostained brain of 201Y>ChT^RNAi^ flies dissected out from the pupal case. (F) shows anti-ChT immunostained brain of 201Y/ ChT^RNAi^;UAS-ChT/UAS-ChT flies rescued by overexpression of *ChT.* (G) Bar graphs showing quantification of E-F, anti-ChT fluorescence signal inside MB α/β lobes (I_α/β_) normalized to ChT signal in neuropilar areas outside MB lobes (I_ON_) in indicated genotypes. For rescue dissected adult brain from 201Y/ ChT^RNAi^;UAS-ChT/UAS-ChT was analyzed. (H) shows percent eclosion in 201Y/ ChT^RNAi^;UAS-ChT and 201Y/ ChT^RNAi^; UAS-ChT/UAS-ChT flies as compared to 201Y>ChT^RNAi^ genotypes. (I) shows mCD8GFP (green) driven by MB247GAL4, marks α/βs and α/βp neurons and co-stained by anti-ChT (magenta) in MB lobe. (J) shows anti-ChT immunostained brain of MB247>ChT^RNAi^ (K) shows anti-ChT fluorescence signal inside MB α/β lobes (I_α/β_) normalized to ChT signal in neuropilar areas outside MB lobes (I_ON_) and (L) shows percent eclosion in MB247>ChT^RNAi^ compared to MB247>+. All images are z-stack pseudocolored representative of 5-7 adult brains. Error bars represent mean ± SEM, *** represent p<0.001. Statistical analysis is based on one-way ANOVA for pairwise comparison. Scale bar, 50 μm

**Table 1:**
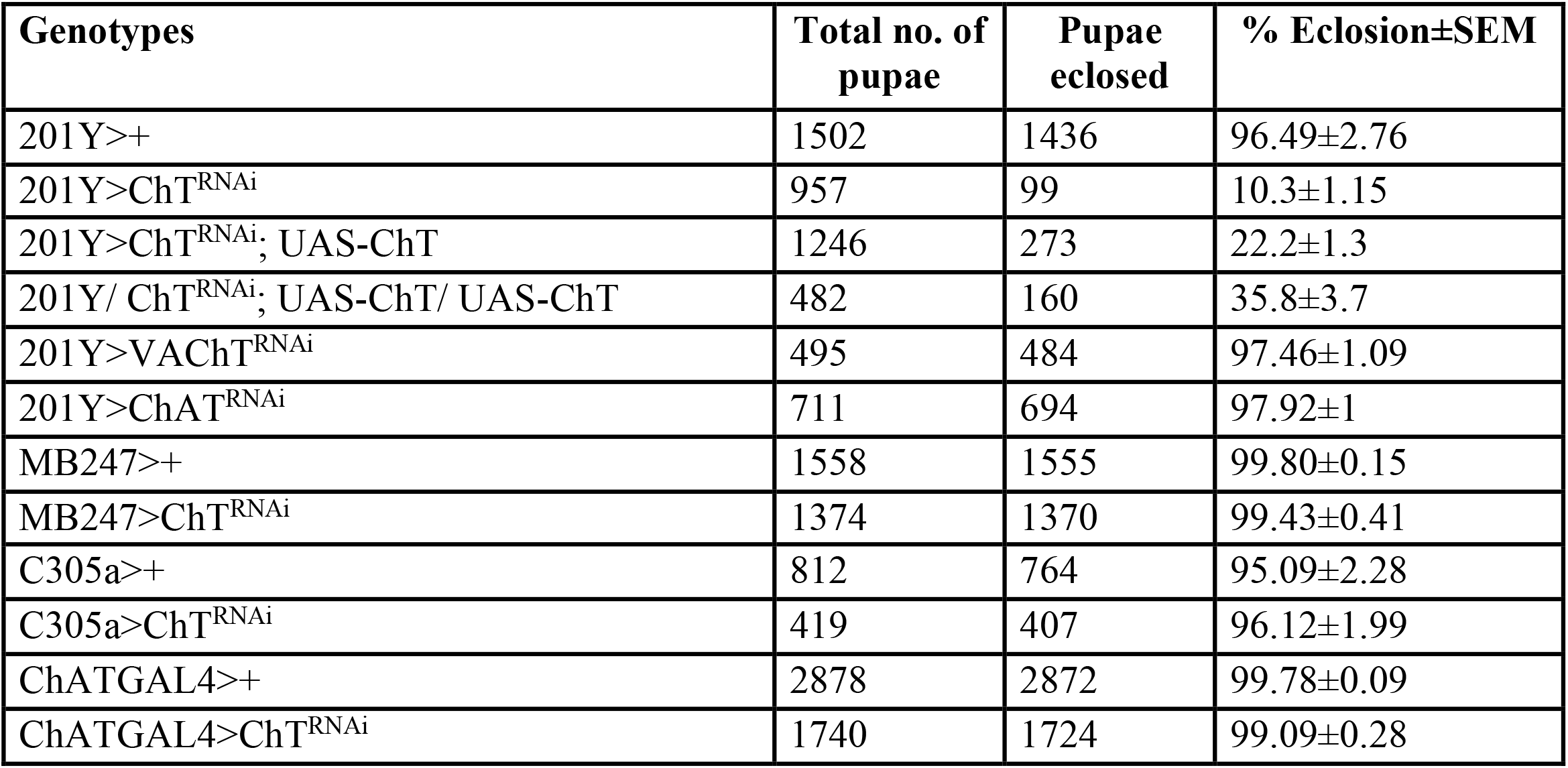
Table summarising a total number of pupae analyzed in this study for eclosion failure due to knock down of ChT in a different subset of Kenyon cells and mushroom body lobes

Since ChT is present ubiquitously in MB, we explored if knockdown of ChT in other lobes of MB also shows similar eclosion failure. We first checked the expression of another GAL4, *MB247GAL4,* using UASmCD8GFP. The progeny showed strongly labeled α/β surface (α/βs), α/β posterior (α/βp) but a very weak expression in α/βc neurons and ϒ lobe (Fig. 2I). Knockdown of ChT by expression of UASChT^RNAi^ in these neurons showed significant reduction of ChT immunoreactivity in the adult brain of MB247GAL4>ChT^RNAi^ (Control 1.02±0.09, N=9; MB247/ChT^RNAi^ 0.58±0.04, N=12, Fig.2 J-K). However, there was no eclosion failure observed (Fig. 2L). Since ChT knockdown in α/βs and α/βp neurons did not show any eclosion defects, we assessed for mobility defects in these flies using climbing assay. Flies are negatively geotactic and have a natural tendency to move against gravity when agitated. We did not find any significant climbing defects due to reduction of ChT in α/βs and α/βp neurons as compared to controls (Fig. S3 A). We next assessed downregulation of ChT in α’/β’ lobe by *C305aGAL4* which also did not show any eclosion failure compared to their genetic controls (Fig. S3 B and Table 1).

Together, these data suggest that ChT in α/βc neurons play a specific role in eclosion of fly from the pupal case. ChT seems to play, as yet unidentified function in other neurons of MB. These findings also support the idea that ChT may have a differential function in different intrinsic neurons of KC.

### The function of ChT in α/βc neurons is independent of the cholinergic pathway

An earlier study using immunolocalization demonstrated that ChAT and VAChT are absent from MB (Gorczyca and Hall, 1987). On the other hand, microarray analysis in another study suggested that ChAT and VAChT are present to comparable amounts both inside and outside of MB lobes (Perrat et al., 2013). More recent study shows that KC expresses ChAT and VAChT and suggest that the majority of KC are cholinergic (Barnstedt et al., 2016). In the present study, we reinvestigated for the presence of ChAT and VAChT along with ChT by immunostainings. We observed that ChT shows immense immunoreactivity in all the lobes of MB whereas ChAT immunoreactivity is not detectable (Fig. S4). Although, VAChT shows little immunoreactivity but from these experiments, we cannot rule out the possibility if it is due to extrinsic neuropilar area. To clarify if there are any cholinergic innervations in MB, we expressed mCD8GFP using *ChAT*GAL4 and co-stained *ChAT*GAL4>ChT^RNAi^ adult brain with anti-ChT. We observed very less amount of mCD8-GFP fibers restricted only to peripheral areas of α/β lobe (Fig. 3A). However, there is a possibility that these fibers belong to the extrinsic cholinergic processes and not to MB. Taken together, our results suggest that either there are very less number of cholinergic innervations in MB or they are altogether absent.

**Figure 3:**
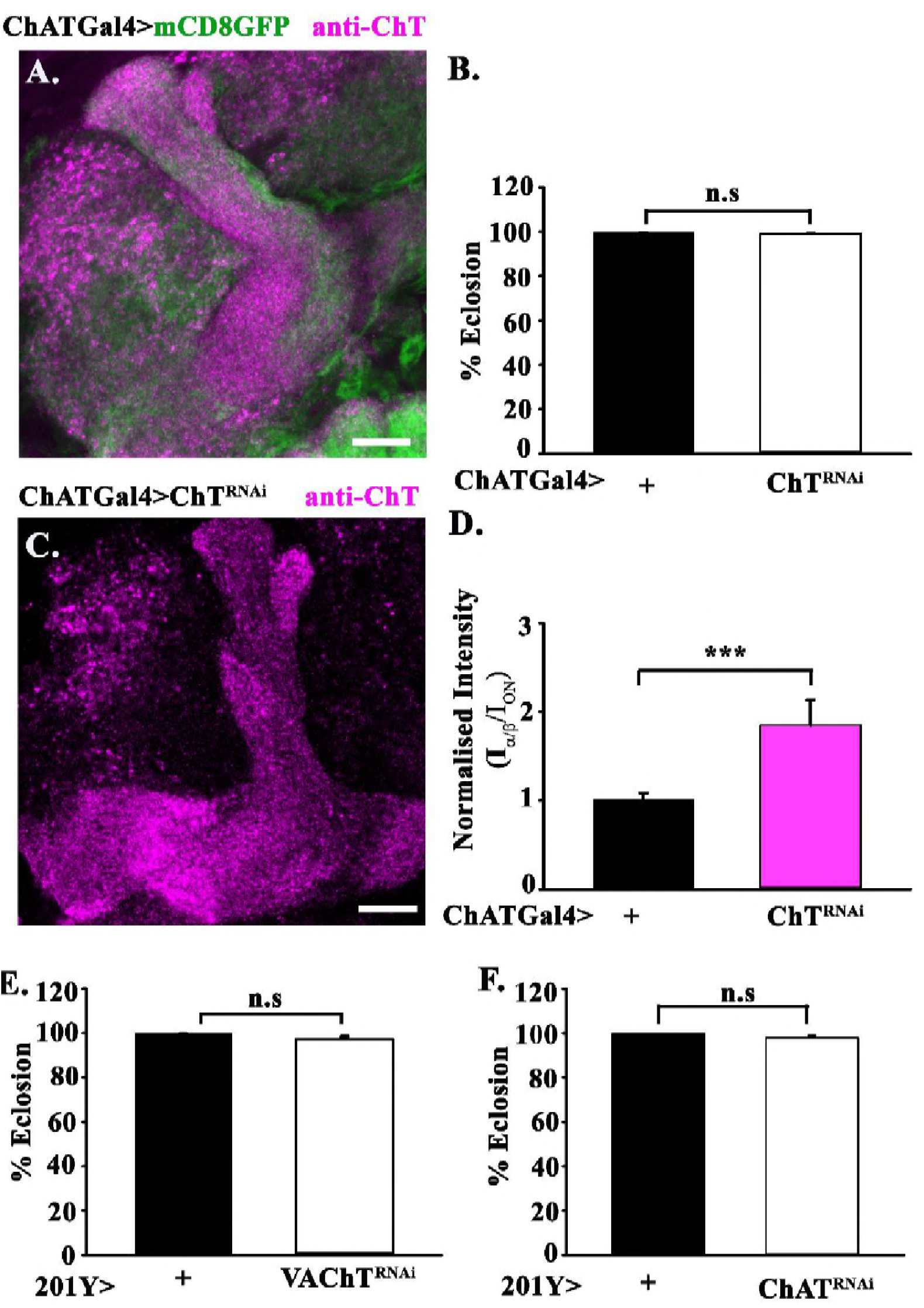
Knockdown of ChT in cholinergic neurons and knock down of cholinergic locus proteins in α/βc neurons do not cause eclosion failure. (A) Shows mCD8GFP driven by ChATGAL4 that marks cholinergic neuropile (green) and co-stained by anti-ChT (magenta) in the MB lobe. (B) shows percent eclosion failure by knockdown of ChT in ChATGAL4>ChT^RNAi^ as compared to ChATGAL4>+. (C) shows anti-ChT immunostained brain of ChATGAL4>ChT^RNAi^ flies and (D) are bar graphs showing quantification of anti-ChT fluorescence signal inside MB α/β lobes (I_α/β_) normalized to ChT signal in neuropilar areas outside MB lobes (I_ON_) in indicated genotypes. (E and F) are percent eclosion by knockdown of ChAT and VAChT in 201Y>ChAT^RNAi^ and 201Y>VAChT^RNAi^ as compared to 201Y>+. All images are a pseudocolored representative image of the 3-5 adult brain. Error bars represent mean ± SEM, *** represent p<0.001. Statistical analysis is based on one-way ANoVA for pairwise comparison. Scale bar, 50 μm

We also investigated if the eclosion failure observed due to depletion of ChT is dependent on cholinergic metabolic cycle or not by downregulating ChT in *ChAT*GAL4>ChT^RNAi^ flies and observed that eclosion was perfectly normal (Fig.3B, Table 1). We quantitated the ChT intensity in *ChAT*GAL4>ChT^RNAi^ adult brain and observed a significant reduction of ChT in outer neuropilar areas (I_ON_) as compared to ChT intensity inside α/β lobes (I_α/β_). The normalised intensity (I_α/β_/I_ON_) in *ChAT*GAL4>ChT^RNAi^ was higher than its genetic control *ChAT*GAL4>+ (Control 1.01±0.08, N=9; *ChAT*GAL4/ChT^RNAi^ 1.89±0.29, N=9, Fig. 3 C-D). To further confirm the non-cholinergic role of ChT in α/βc MB neurons in eclosion, we downregulated VAChT^RNAi^ and *ChAT*^RNAi^ in α/βc neurons and did not observe any eclosion failure as compared to its genetic control (Fig. 3 E-F). Taken together, these data strongly suggest that pupal eclosion failure is specifically due to ChT downregulation in α/βc neurons. The function of ChT in these neurons are functionally uncoupled from cholinergic locus suggesting α/βc neurons being non-cholinergic in nature. It also indicates that ChT is present in a much larger number of KCs which might necessarily not be cholinergic.

### ChT function in α/β core neurons regulate peristalsis through a cholinergic independent pathway

Eclosion of flies from the pupal case involves coordinated contraction and relaxation of whole body muscles (Kimura and Truman, 1990; Rivlin et al., 2004). This process helps in the forward movement to drive the fly out of the pupal case. To understand the eclosion failure, we assessed the peristaltic movement of late 3^rd^ instar *Drosophila* larvae from caudal to rostral side (forward movement) in 201YGAL4>ChT^RNAi^ flies. We observed that peristaltic counts were drastically reduced when ChT was depleted in α/βc neurons in 201YGAL4>ChT^RNAi^ (47±0.99 counts/min, N=40) as compared to their genetic controls *201Y*GAL4>+ (61±0.80 contractions/min, N=40) (Fig. 4A). Transgenic over-expression of *ChT* in these neurons restored the peristaltic decrement (62.41±2.02 counts/min, N=40, Fig. 4A). To further clarify if the decrease in the peristaltic count is through the cholinergic mode of action and whether this process involves cholinergic proteins, we downregulated ChAT by expression of ChAT^RNAi^ in α/βc neurons. We did not observe any significant peristaltic decrease in 201YGAL4>ChAT^RNAi^ larvae (57.46±1.25 counts/min, N=40) (Fig. 4A). This suggests that the role of ChT is functionally uncoupled from ChAT in MB α/βc neurons. Locomotion is one of the important behavior of an animal that largely depends on ACh (Rand, 2007). Therefore, we assessed the effect of ChT downregulation on peristalsis in cholinergic neurons also. Downregulation of *ChT* in cholinergic neurons reduces peristaltic count (43.46±2.42 counts/min, N=40) as compared to their genetic control (60±1.82 counts/min, N=40) (Fig. 4B). This decrement in peristalsis is normalized by transgenic over-expression of *ChT* in cholinergic neurons (61±1.91contractions/min, N=40) (Fig. 4B). Intriguingly, like in α/βc neurons, depletion of *ChAT* by ChAT^RNAi^ in cholinergic neurons also does not reduce peristaltic counts (60.13±0.89 counts/min) as compared to their genetic control (Fig. 4B). However, we observed a developmental delay by 3-4 days at 29 ^o^C on knockdown of ChAT in cholinergic neurons as compared to their genetic controls. Taken together, these observations suggest that ChT can have cellular roles independent of the canonical cholinergic pathway.

**Figure 4:**
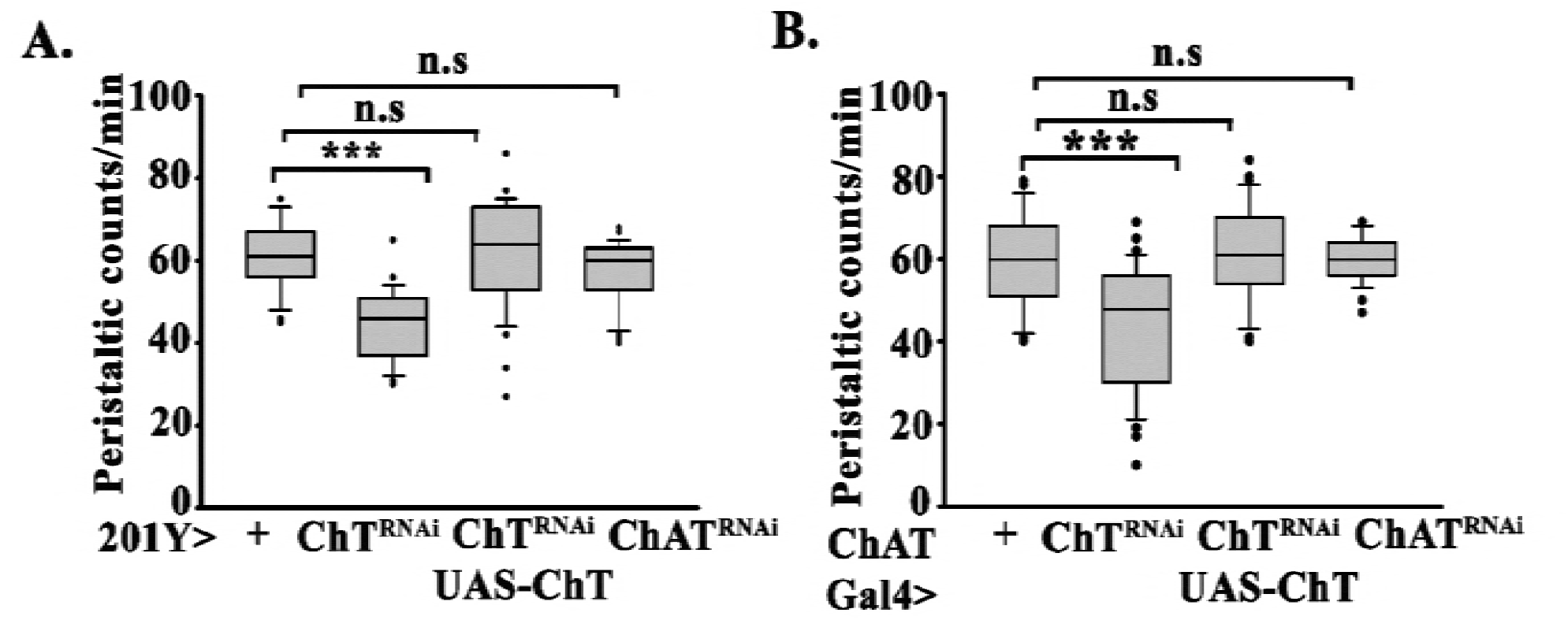
Knockdown of ChT in α/βc neurons and cholinergic neurons alter the peristaltic behavior of 3^rd^ instar larvae. A and B show peristalsis quantified as a number of body wall contractions from posterior to anterior end and represented as peristaltic counts per min in the indicated genotypes. Distribution of data is shown as a box plot (n=40) for each genotype. The box plot show box boundaries as 25% and 75% quartiles and median as the center line. 10% and 90% quartiles are shown as whiskers. *** represent p<0.001 and n.s represent nonsignificance. Statistical analysis is based on one-way ANOVA for pairwise comparison. Kruskal Wallis oneway analysis on variance on ranks was done where the normality test fails.

### ChT function in MB α/β core neurons contributes to NMJ maintenance

Proteins like *Drosophila* neurexins (DNRx), Scribble and RanBPM have been shown to be present in MB and are also associated with NMJ phenotype (Rui et al., 2017; Scantlebury et al., 2010). Therefore, we determined if the locomotor defect caused by downregulation of ChT in α/βc neurons could be due to alteration in NMJ morphology. We knocked down *ChT* in α/βc neurons and assessed NMJ morphology in 3^rd^ Instar larvae. We found a significant increase in the number of boutons per muscle area at the NMJs of *201Y*>ChT^RNAi^ (1.65 ± 0.06, N=16) as compared to their genetic control, *201Y*>+ (1.32±0.03, N=16) (Fig. 5 A-B and F). There was also an increase in the number of branches in *201Y*>ChT^RNAi^ (10±0.85, N=16) as compared to the genetic control (5±0.57, N=16) (Fig. 5 A-B and G). This phenotype was significantly rescued by transgenic over-expression of *ChT* in α/βc neurons (Boutons, 1.34±0.05, N=10; no. of branches, 8.5±0.42, N=11) (Fig.5 C and H-I). Interestingly, downregulation of ChT in cholinergic neurons of ChATGAL4>ChT^RNAi^ larvae did not show any significant alteration in bouton number (1.09±0.07, N=10) as well as a number of branches (8.5±0.83, N=10) as compared to the controls (Boutons, 1.05±0.06; no. of branches, 6.5±0.37) (Fig. D-G). Therefore, we propose that peristaltic count decrement due to ChT knockdown in α/βc neurons may be due to improper NMJ functioning that involves a non-cholinergic pathway for its maintenance. Its decrement in cholinergic neurons affecting peristalsis could be through a different mode of action that involves cholinergic pathway. However, the mechanisms by which ChT controls larval peristaltic movement through a pathway that does not require ChAT in MB α/βc as well as in cholinergic neurons requires further investigation. Together these results further corroborate our observations that ChT function in α/βc neurons is independent of the cholinergic pathway.

**Figure 5:**
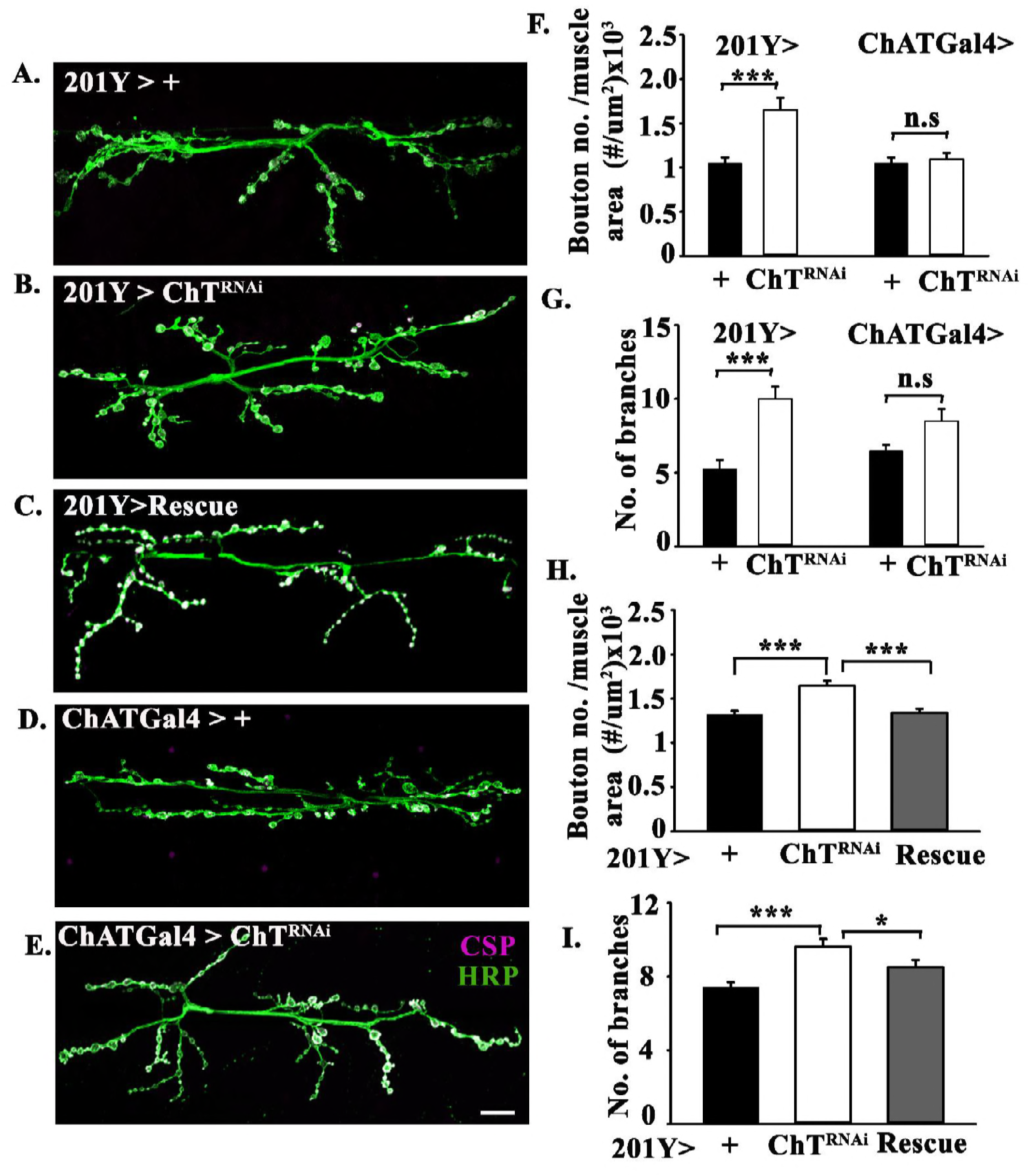
Knockdown of ChT in α/βc neurons but not in cholinergic neurons alter NMJ phenotype in 3^rd^ instar larvae. Representative images of the given genotypes at muscle 6/7 of A2 hemisegment in 3^rd^ instar larva; (A) 201Y>+, (B) 201Y>ChT^RNAi^, (C) 201Y/ChT^RNAi^; UAS-ChT/UAS-ChT (Rescue), (D) ChATGal4>+, (E) ChATGAL4>ChT^RNAi^. The NMJs shown were stained with anti-HRP (green) and anti-CSP (magenta). Scale bar in E represents 20 μm. (F-I) Bar graphs showing number of boutons and the total number of branches per muscle area of 6/7 muscle of A2 hemisegment. Error bars represent mean ± SEM; N= 12-17; *** represent p< 0.001; * represent p<0.05. Statistical analysis was done using the two-tailed t-test and non-parametric Mann-Whitney rank sum test where the normality test failed.

## Discussion

In the present study, we report that ChT is ubiquitously present in MB and its presence in a specific subset of MB neurons called α/βc neurons is important for eclosion from pupae of *D. melanogaster,* locomotory behavior, and maintenance of NMJ integrity. Here, we present evidence that the role of ChT in MB α/βc neurons is functionally uncoupled from the cholinergic locus. Together our results suggest that ChT can affect different downstream functional pathways which can be either cholinergic or non-cholinergic. Our findings thus establish a new avenue for ChT study, that, it is merely not an integral protein of the cholinergic cycle but also has other potential biological functions.

α/βc neurons are the last formed subset of neurons of α/β lobes which are formed between the late pupal stage until adult eclosion (Strausfeld et al., 2003). Our observations that pupal lethality was seen only when ChT was downregulated in α/βc but not in α/βp, α/βs neurons of α/β lobe or in α’/β’ lobe suggests that ChT plays distinct functional roles in different subsets of MB neurons. We do not observe any defect in the organization of the MB lobes on downregulation of ChT suggesting that it does not have a role in the organization of axonal fibers in these lobes. It is worth mentioning here that ChT knock out mice also showed neonatal lethality while the gross organ development and morphology of the pups were normal (Ferguson et al., 2004). This is closely similar to our observations of eclosion failure and consequent pupal lethality in *Drosophila* due to knock down of ChT. One possible cause of eclosion failure may be the altered NMJs that are observed due to knock down of ChT. The underlying mechanism of how ChT in α/βc neurons affects NMJ morphology is currently unclear. However, like ChT, several other proteins, namely; *Drosophila* Neurexins, Scribble, adaptor protein DRK and Wallenda have also been reported to have localisation in MB and play a role in maintaining structural plasticity at NMJs via different signalling cascades (Moressis et al., 2009; Rui et al., 2017; Shin and DiAntonio, 2011). Previous reports also describe that MB physiology regulates locomotor activity rhythms in *Drosophila* (Gorostiza et al., 2014; Mabuchi et al., 2016). Future studies are required to elucidate the potential downstream circuit that links the role of ChT in MB physiology and MB motor output.

For an efficient cholinergic neurotransmission, all the components of the ACh metabolic cycle should be present at the synaptic junctions. While there is a predominant expression of ChT in MB, our immunostaining analysis shows that ChAT and VAChT are either absent or present in a negligible amount in MB which is undetectable at endogenous levels. In the current study, we provide multiple evidences that support non-canonical functions of ChT in α/βc neurons: the downregulation of ChT but not ChAT or VAChT in α/βc neurons of MB causes eclosion failure suggesting that ChT regulate eclosion through a pathway that is functionally uncoupled from the cholinergic locus. Vice-versa, ChT knockdown in cholinergic neurons does not produce eclosion failure suggesting that α/βc neurons are non-cholinergic. These observations corroborate the idea that ChT can have a non-canonical functional role at least in α/βc neurons. Our assertion of the non-canonical role of ChT in MB is further supported by our observations that ChT knockdown in α/βc neurons leads to altered phenotype at NMJs showing increased boutons and branch number. This phenotype was not observed when ChT was downregulated in cholinergic neurons. Furthermore, we see a reduction of peristaltic count on knockdown of ChT but not by ChAT in α/βc neurons. Indeed, in NSC-19 cells expression of cholinergic locus and ChT was reported to be differentially regulated (Brock et al., 2007). Although we did not detect ChAT and VAChT in MB but our data do not rule out any non-cell autonomous function of ACh affecting α/β lobe functioning.

Alternatively, ChT in MB may regulate different functions through an indirect downstream pathway. Chromatin remodelers like histone acetyltransferase (HAT) and Histone deacetylase (HDAC4) are dependent on the levels of choline (Ward et al., 2013). They also act as regulators of transcription factors like *D-Mef2* in *Drosophila* MB (Fitzsimons et al., 2013; Fogg et al., 2014) and *FOXP3* in mammals (Li and Greene, 2007). *FOXP* is present in MB and has been associated with locomotor defects and memory deficits (DasGupta et al., 2014). Indeed, we observed similar eclosion failure when we knocked down ChT with Mef2GAL4 (data not shown). In the context of our observations, it is possible that ChT in α/βc neurons is required for the uptake of choline into these neurons, not for ACh synthesis but different regulatory pathway involved in developmental and behavioral processes. This could be the possible reason that we see eclosion failure and locomotory defects due to ChT knockdown but not by reduction of cholinergic proteins. Thus, we propose that ChT maintains required levels of choline in α/βc neurons for different downstream processes other than ACh synthesis.

Our findings have important implication in redefining the biological role of ChT to affect downstream pathway which may not be cholinergic. We speculate that ChT may be present in the cell types that require a high amount of choline and not just cells that synthesize ACh. Thus, we anticipate that the role of ChT is much far-reaching than previously thought. Although neuroanatomy of flies and vertebrates are quite distinct but the proteins of cholinergic signaling pathway are highly conserved. We believe that it will encourage further investigation into the developmental role of ChT as well as the role it plays in learning and memory in both invertebrates and higher organisms.

## Material and methods

### Drosophila stocks and culture conditions

All *Drosophila* stocks and their crosses were grown on standard cornmeal/agar media supplemented with yeast at 25 ^o^C, under a 12-12 hr light-dark cycle. All crosses for RNAi experiments were grown at 29 ^o^C. For all control experiments, GAL4 drivers were crossed with w^1118^ (+) were used, unless otherwise mentioned in the experiments. The UASRNAi strains for ChT (101485), VAChT (32848) were obtained from Vienna *Drosophila* RNAi Center (VDRC), Vienna, Austria and for ChAT (25856) was obtained from Bloomington stock center, Bloomington, Indiana. The GAL4 drivers used were *Elav^C755^GAL4* (458), ChATGAL4 (6798), *201Y*GAL4 (4440), *MB247GAL4* (50742), *c305aGAL4* (30829).

### Cloning and generation of anti-ChT polyclonal antibody

cDNA fragment corresponding to the hydrophilic ChT C-terminal domain (Glu-489 to Phe-614) was amplified by PCR using two oligonucleotide primers 5’AAGGATCCATGGAGTCCGGCAAGTTGCCGCCCA3’ and 5’AAAAGCTTTCAGAAGGCCGTATTGTCCT 3’. The amplified fragment was inserted into the BamHI/HindIII site of the pGEX-KG fusion protein expression vector. The fusion protein was purified and the protein domain was later cleaved from the glutathione S-transferase by incubation with thrombin overnight at 10°C followed by SDS-polyacrylamide gel electrophoresis to assess the extent of cleavage. The cleaved fragment was eluted and used for immunizing rabbits. About 250 μg of protein was used for the first immunization. For each booster doses (6x) 100 μg was used (Deshpande laboratories, Bhopal, India). The antibody from serum was later affinity purified before use.

### Generation of UAS-ChT transgenic line

The open reading frame of ChT was amplified from cDNA using a gene-specific forward primer (5’AAGAATTCATGATCAATATCGCTGGCG-3’) and reverse primer (5?AGCGGCCGCTCAGAAGGCCGTATTGTCCT3’). The amplified fragment was cloned between EcoRI and NotI site of the pUASt vector and injected into *Drosophila* embryos for transgenesis.

### Antibodies and immunohistochemistry of larval and adult brain

The primary antibodies used were: rabbit anti-ChT (1:300, this study), mouse anti-ChAT (1:1000, 4B1, DSHB), mouse anti-CSP (1:20, DSHB), anti-VAChT (1:200, a gift from Toshihiro Kitamoto, U. Iowa, Iowa City, IA). Conjugated secondary antibodies used were Alexa Fluor-568 (Molecular Probes), Alexa Fluor 647 (Jackson ImmunoResearch).

For immunohistochemistry, third instar larvae were age-matched and dissected in PBS as previously described (Baqri et al., 2006). For the adult brain, flies were anesthetized on ice and brain tissue was dissected out in cold PBS. Subsequently, tissues were fixed with freshly prepared 4% paraformaldehyde for 1.5 hr, washed and incubated with primary antibody at 4°C overnight. Following day, the tissues were incubated with secondary antibodies for 1.5 hr at room temperature, washed and finally mounted in vecatashield in between the bridge prepared by double sided tape in order to avoid squashing of tissue directly under the coverslip. All images were collected using Leica SP8 LSCM using oil immersion 63x/1.4 N.A objective and subsequently processed using Image J 1.50i (NIH, USA). Pupal images were taken using Zeiss Axiocam ERC 5S mounted on Zeiss Stemi2000 CS stereomicroscope.

For antibody quantification, three rectangular ROI of 50 X 50-pixel size was drawn over the α/β lobe. The mean fluorescence in the three ROI in α/β lobe (I_α/β_) was calculated using Image J. The α/β lobe intensity was normalized with respect to mean fluorescence intensity of the three ROI of 50 X 50-pixel size in the neuropilar area outside MB (I_ON_).

### Estimation of eclosion failure

Crosses were set between 4-5 days old males and virgin females in the ratio of 4:8 and left overnight at 29°C. Following day, the first vial was discarded and flies were subsequently transferred to new ones with fresh media, every 24 hours, for the next seven days. For each day number of total pupae developed and empty pupal cases were counted. Pupal lethality was scored on the basis of percentage eclosion calculated by [(Number of empty pupae/Total number of pupae) X100]. The quantitative and statistical analysis was performed in Sigma Plot ver. 12.5. One-way ANOVA followed by *post-hoc* Tukey test for pairwise comparison was used.

### Peristalsis assay

Peristalsis assay was done manually on 15 cm petridish containing 2% hardened agar. Larvae were washed and kept on the plate for 1 min acclimatization. Subsequently, the peristaltic contractions were counted for 1 min under a dissection microscope. Full posterior to the anterior movement was counted as one contraction. The larvae which did not move were not considered in the analysis. The quantitative and statistical analysis was performed in Sigma Plot ver. 12.5. One-way ANOVA followed by *post-hoc* Tukey test for pairwise comparison was used.

### Climbing assay

To determine the climbing activity of the flies, the assay vial was prepared by vertically joining two empty polystyrene vials with open ends facing each other. The vertical distance was marked at 6 cm above the bottom surface. A group of 10 flies irrespective of the gender was transferred to fresh vials for 24 hours. For the assay, flies were transferred into the assay vial and allowed to acclimatize for 30 minutes. The flies were then gently tapped down to the bottom of the vial and the number of flies that climb above the 6 cm mark was counted. For each vial three trial readings were taken, allowing for 1 min rest in between each trial. A total of 12 groups were assayed for each genotype. The quantitative and statistical analysis was performed in Sigma Plot ver. 12.5. One-way ANOVA followed by *post-hoc* Tukey test for pairwise comparison was used.

### Immunohistochemistry and morphological quantification of NMJs

Third instar larvae were dissected in the calcium^-^free HL-3 buffer and fixed in 4% paraformaldehyde for 30 min. Subsequently, larvae were washed with 1X PBS, 0.2% Triton X-100 and blocked in 5% BSA for 1 hour. Incubation of larval fillets was done overnight at 4^0^C in primary antibody and then with secondary antibodies for one and half hour at room temperature and mounted in Flourmount G (Southern Biotech). Antibodies used were mouse anti-CSP (ab49-DSHB) in 1:50 and Alexa 488 conjugated anti-HRP in 1:800 (Molecular Probes). Species-specific fluorophore conjugated secondary antibody used was Alexa 568 in 1:800 dilution.

Morphological quantification of NMJ was performed at muscle 6/7 in the A2 hemisegment. For quantification of different parameters of NMJ, morphology images were captured at 40X objective in Olympus FV3000. Image J (NIH) and cellSens software were used for muscle area and Bouton number. Bouton number was normalized to respective muscle area. A total number of branches were quantified as described earlier (Coyle et al., 2004). Statistical analysis was done by using students t-test and non-parametric Mann-Whitney rank sum test where the normality test failed. All data values indicate mean and standard error mean.

## Acknowledgements

We acknowledge Department of Science & Technology, Govt. of India for financial support to R.H vide reference no. SR/WOS-A/LS-77/2013 under Women Scientist Scheme-A to carry out this work. We acknowledge Indian Institute of Science Education and Research, Bhopal (IISER-B) for hosting R.H during initial tenure of the project. We thank Toshihiro Kitamoto, for providing *anti-VAChT* antibody, Bloomington Drosophila Stock Center (BDSC), Indiana USA and Vienna Drosophila RNAi Centre (VDRC), Austria for fly stocks. We also thank Krishanu Ray and Manish Jaiswal for many useful comments on the manuscript.

## Author contribution

R.H conceived the project, designed and performed most of the experiments. N.H performed eclosion estimation, climbing assays with assistance from R.H. S.K performed NMJ experiments. S.S. performed peristalsis assay. R.H analysed and interpreted the results with critical inputs from N.H. R.H compiled the figures and wrote the paper. V.K provided resources for antibody and transgenic flies generation. V.K and R.K.M gave scientific inputs, editing, critical comments and provided resources.

## Reagent availability

All the reagents generated and used in the manuscript will be shared with the scientific community.

## Supplementary figures

**Figure S1:**
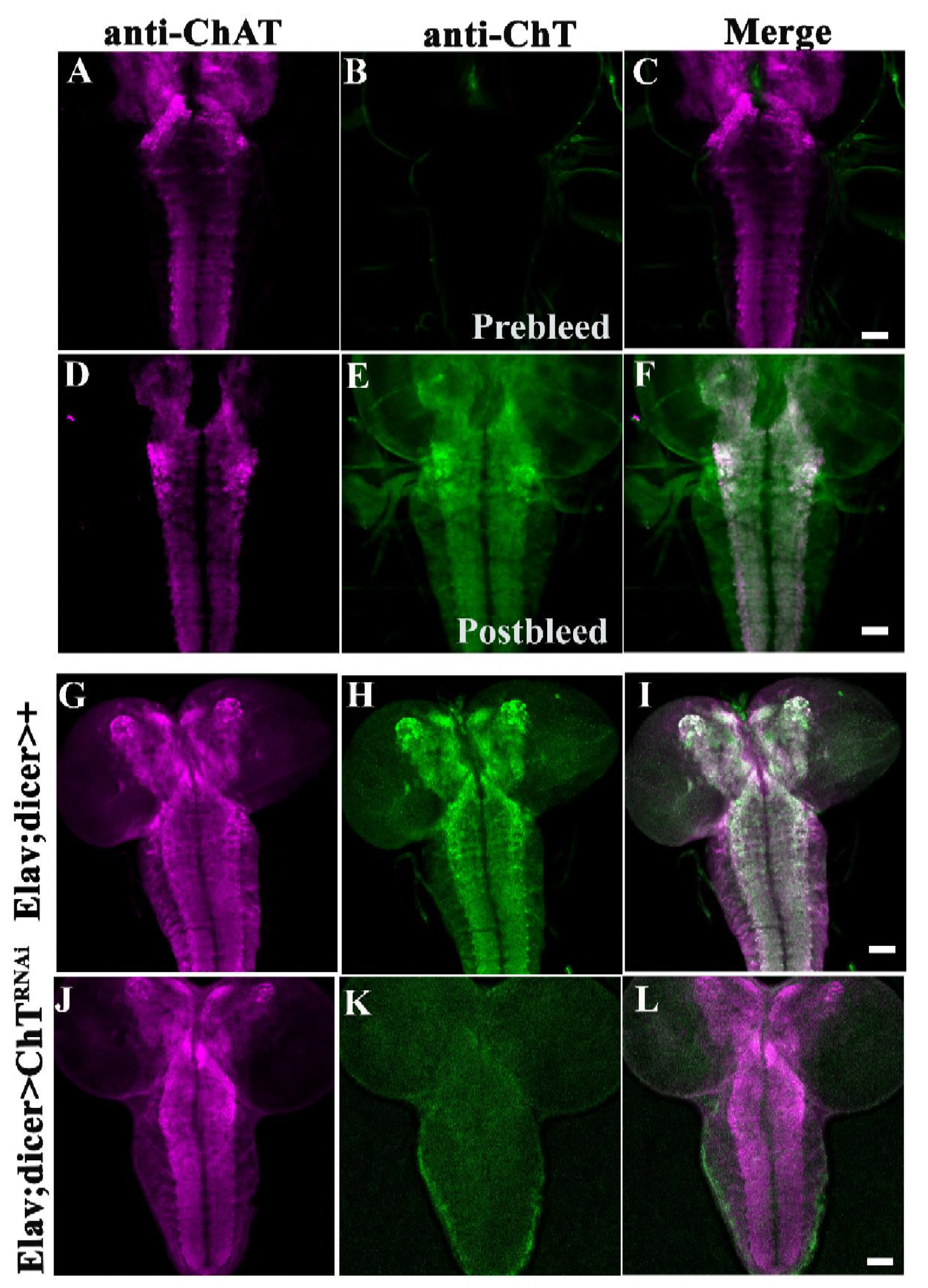
Drosophila polyclonal anti-ChT antibody specifically detects endogenous ChT proteins. (A) Immunostaining with *anti-ChAT* (4B1) antibody (magenta), (B) pre-bleed serum from rabbit, (C) merge image of A and B, (D) showing endogenous ChAT localization with anti-ChAT at neuropil of 3^rd^ instar larval brain, (E) immunostaining with post-bleed serum (affinity purified) (green) (F) merged image showing colocalization of ChAT and ChT. (G-L) shows drastic reduction of ChT protein in Elav;;dicer>ChT^RNAi^ compared to Elav;;dicer>+ (green). Co-immunostaining with anti-ChAT was used as a control (magenta). All images are pseudocolored z-stack confocal images of third instar larval VNC, a representative image of immunostaining done in the 3-5 brains in each case. Scale bar, 100μm.

**Figure S2:**
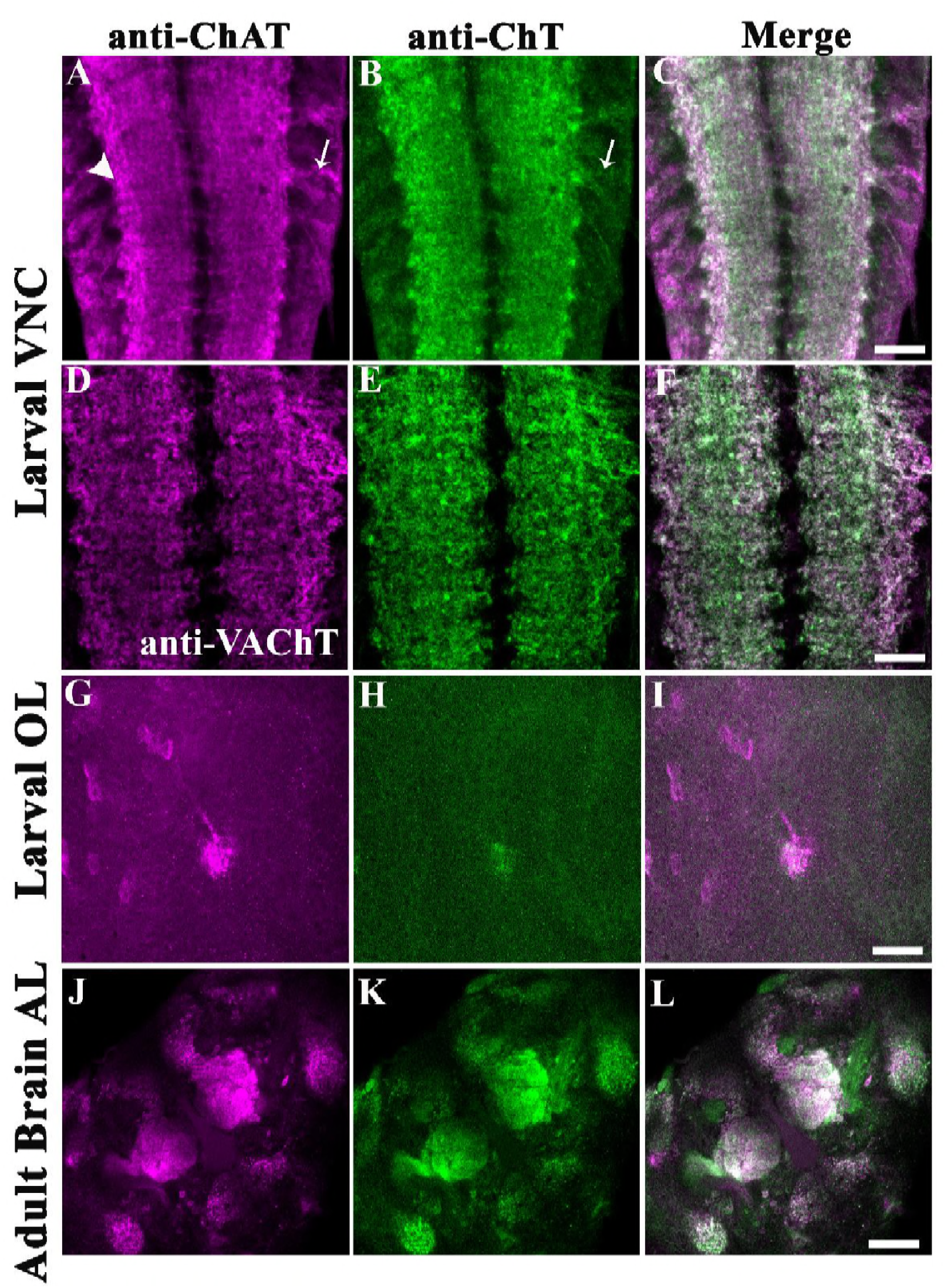
ChT colocalizes with markers of cholinergic neurons and in areas with predominant cholinergic synapses. (A-C) Co-immunostaining with anti-ChAT (magenta), anti-ChT (green), and colocalization of ChAT and ChT as a merged image. (D-F) with anti-VAChT (magenta), anti-ChT (green), and colocalization of VAChT and ChT as a merged image. (G-I) shows colocalization of ChAT and ChT in Bolwig nerve in larval optic lobes (OL). (J-L) shows colocalization of ChAT and ChT in antennal lobes (AL) of the adult fly brain. Images shown here are the representative image of immunostaining done in the 3-5 brains in each case. Scale bar, 50μm

**Figure S3:**
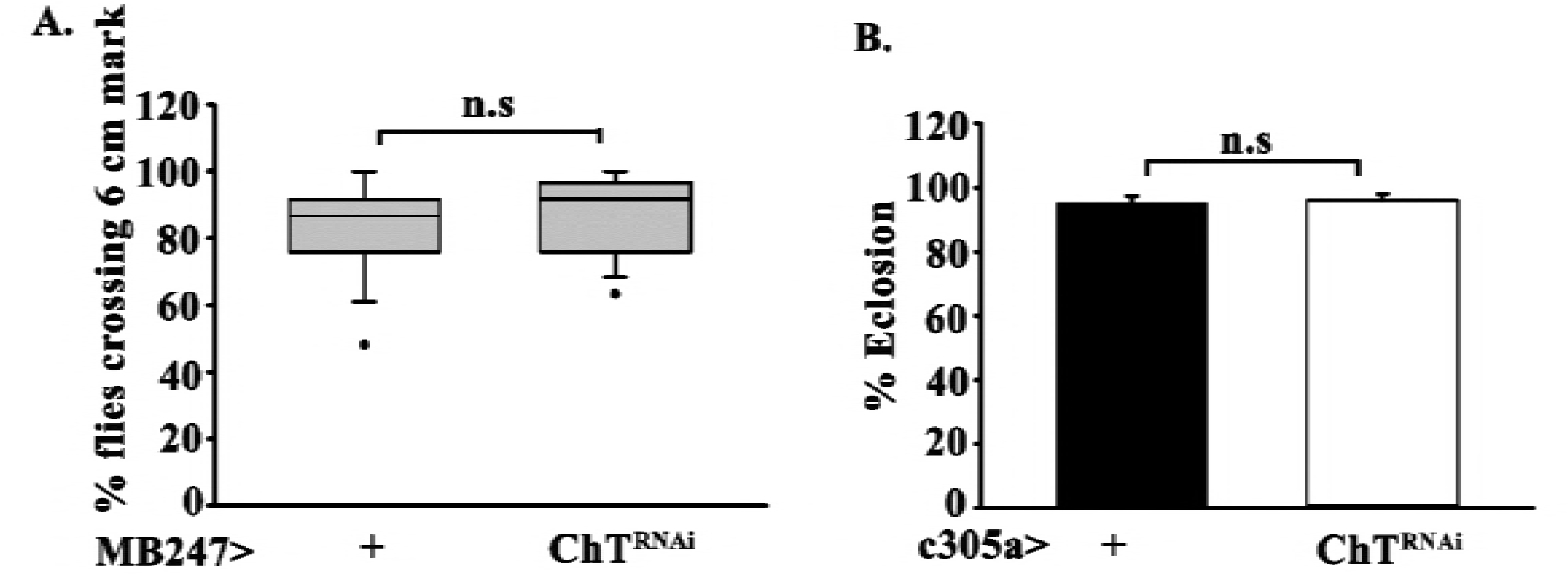
(A) Box plot showing percent climbing activity of MB247>ChT^RNAl^ flies crossing 6 cm mark per 10 secs as compared to MB247> +. Data shown here is derived from 12 assay vials per genetic group. Each assay vial containing a mixed population of 10 male and female flies, n.s denote non-significance. (B) percentage eclosion of c305aGAL4>ChT^RNAi^ pupae as compared to its genetic control c305a>+. n.s denote nonsignificance. Statistical significance was calculated using one-way ANOVA by *post-hoc* Tukey test for pairwise comparison.

**Figure S4:**
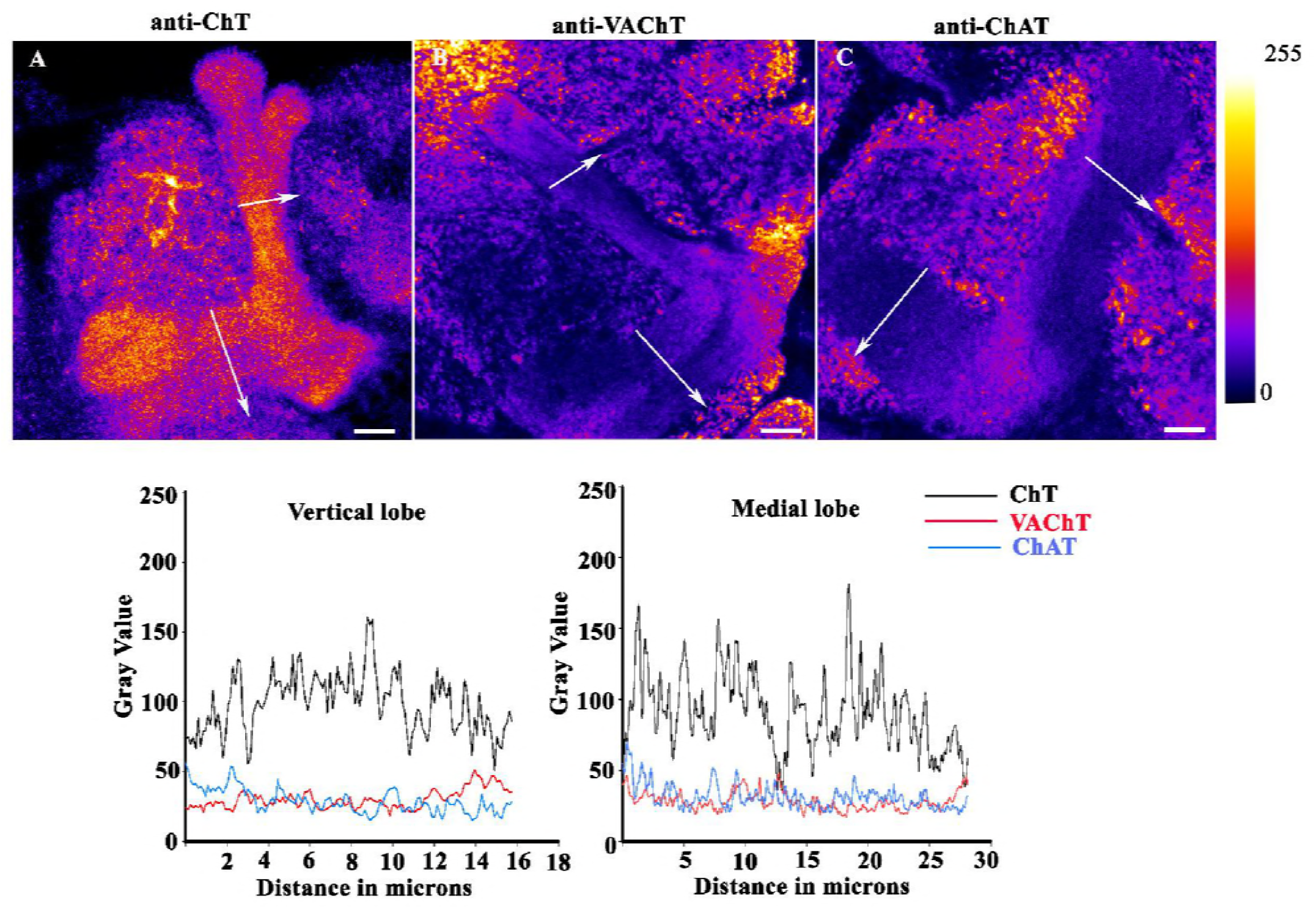
Endogenous ChT is predominantly expressed in MB lobes as compared to its canonical proteins of the cholinergic locus, ChAT, and VAChT. (A-C) show images of MB lobes immunostained with anti-ChT (A), anti-VAChT (B) and anti-ChAT (C) converted to Fire LUT map (using image J) that shows the scale of colors from 0-pixel intensity (minimum) till 255-pixel intensity (maximum). White arrows show the direction of line intensity plot whose absolute values are shown for vertical lobe (D) and Medial lobe (E) for ChT (black), VAChT (red), ChAT (blue). Scale bar, 50μm

**Video 1:** Video showing 201Y driven ChT^RNAi^ pupae are alive till more than the P14 stage of pupal development.

**Video 2:** Video showing 201Y driven ChT^RNAi^ fly which partially ecloses and gets stuck in the pupal case.

